# Znf598-mediated Rps10/eS10 ubiquitination contributes to the ribosome ubiquitination dynamics during zebrafish development

**DOI:** 10.1101/2023.02.12.528238

**Authors:** Nozomi Ugajin, Koshi Imami, Hiraku Takada, Yasushi Ishihama, Shinobu Chiba, Yuichiro Mishima

## Abstract

Ribosome is a translational apparatus that comprises about 80 ribosomal proteins and four rRNAs. Recent studies reported that ribosome ubiquitination is crucial for translational regulation and ribosome-associated quality control (RQC). However, little is known about the dynamics of ribosome ubiquitination under complex biological processes of multicellular organisms. To explore ribosome ubiquitination during animal development, we generated a zebrafish strain that expresses a FLAG-tagged ribosomal protein Rpl36/eL36 from its endogenous locus. We examined ribosome ubiquitination during zebrafish development by combining affinity purification of ribosomes from *rpl36*-FLAG zebrafish embryos with immunoblotting analysis. Our findings showed that ubiquitination of ribosomal proteins dynamically changed as development proceeded. We also showed that during zebrafish development, the ribosome was ubiquitinated by Znf598, an E3 ubiquitin ligase that activates RQC. Ribosomal protein Rps10/eS10 was found to be a key ubiquitinated protein during development. Furthermore, we showed that Rps10/eS10 ubiquitination-site mutations reduced the overall ubiquitination pattern of ribosome. These results demonstrate the complexity and dynamics of ribosome ubiquitination during zebrafish development.

## Introduction

Ribosome is a large ribonucleoprotein complex that plays a central role in translation in all living organisms. The eukaryotic 80S ribosome consists of about 80 ribosomal proteins and four ribosomal RNA (rRNA) molecules organized into the large 60S subunit and the small 40S subunit (Bashan and Yonath 2008; De La Cruz et al. 2015). While there are some species-specific differences, these core ribosomal components are essential constituents of the ribosome to function as a translation apparatus. In addition, post-translational modifications on ribosomal proteins modulate the functionality of the ribosome (Dalla Venezia et al. 2019; Emmott et al. 2019; Li and Wang 2020; Simsek and Barna 2017; Xue and Barna 2012). Ribosomal proteins have been reported to undergo a range of post-translational modifications, including phosphorylation (Imami et al. 2018; Martin et al. 2014), ubiquitination (Back et al. 2019; Higgins et al. 2015; Matsuki et al. 2020; Silva et al. 2015; Spence et al. 2000), methylation (Małecki et al. 2021; Matsuura-Suzuki et al. 2022), acetylation (Zhang et al. 2022), and UFMylation (Walczak et al. 2019; Wang et al. 2020). Some of these modifications are introduced to the ribosome in response to various stimuli, and affect multiple aspects of translation, such as translation efficiency (Matsuura-Suzuki et al. 2022; Takehara et al. 2021), and selectivity (Imami et al. 2018; Matsuki et al. 2020; Zhang et al. 2022). Hence, ribosomes in a given cell or organism are subjected to dynamic changes in their modification status to adapt to diverse translational demands.

Recent studies have emphasized the significance of ribosome ubiquitination in the context of translating ribosome quality control. When a ribosome slows aberrantly during translation, it collides with the trailing ribosome to form a specific structure called as a disome (Ikeuchi et al. 2019; Juszkiewicz et al. 2018). Ribosome collisions and disome formation trigger the ribosome-associated quality control (RQC) pathway, which dissociates stalled ribosomes into subunits and degrades nascent polypeptides (Inada 2020; Joazeiro 2017, 2019). The site-specific ubiquitination of the 40S ribosomal proteins Rps10/eS10 and Rps20/uS10 in the disome initiates the RQC pathway (Garzia et al. 2017; Juszkiewicz and Hegde 2017; Matsuo et al. 2017; Sundaramoorthy et al. 2017). These reactions are catalyzed by an E3 ubiquitin ligase Znf598 (Hel2 in yeast), which recognizes the disome structure by an unknown mechanism (Garzia et al. 2017; Juszkiewicz and Hegde 2017; Juszkiewicz et al. 2018; Matsuo et al. 2017; Sundaramoorthy et al. 2017). Ubiquitinated forms of Rps10/eS10 and/or Rps20/uS10 are recognized by the ASC-1 complex (also known as RQT complex in yeast), in which an ATP-dependent RNA helicase Ascc3 (Slh1/Rqt2 in yeast) dissociates the stalled ribosome into subunits (Juszkiewicz et al. 2020b; Matsuo et al. 2017, 2020; Narita et al. 2022; Sitron et al. 2017). Another E3 ubiquitin ligase, Ltn1, polyubiquitinates a nascent polypeptides in the split 60S subunit, which are then degraded by the proteasome (Bengtson and Joazeiro 2010; Brandman et al. 2012; Shao and Hegde 2014). The ubiquitination also promotes endonucleolytic cleavage of the mRNA with the stalled ribosome, known as no-go decay (NGD) (D’Orazio et al. 2019; Glover et al. 2020; Ikeuchi et al. 2019). In addition to the RQC and NGD pathways, an E3 ubiquitin ligase called Rnf10 (Mag2 in yeast) ubiquitinates the 40S ribosomal proteins Rps2/uS5 and Rps3/uS3 in response to decoding and initiation/elongation abnormalities. This results in the degradation of 40S subunits (Garzia et al. 2021; Li et al. 2022; Sugiyama et al. 2019). Monoubiquitination of Rps3/uS3 by Mag2 is followed by polyubiquitination by Fap1 and Hel2, leading to 18S nonfunctional rRNA decay (18S NRD) (Li et al. 2022; Sugiyama et al. 2019). Overall, these ribosome ubiquitination pathways maintain cellular proteostasis and protect the eukaryotic translation system against a variety of translation elongation issues.

Ribosome ubiquitination is not limited to the quality control system. Rather, several studies revealed that ribosome ubiquitination is linked to a wide range of biological activities. In yeast, at least 37 ribosomal proteins are ubiquitinated in response to oxidative stress (Back et al. 2019). Furthermore, some site-specific ribosome ubiquitinations are required for cells to survive in ER stress (Higgins et al. 2015). Another example is the ubiquitination of the ribosomal protein Rpl28/uL15 during the S phase of the cell cycle (Spence et al. 2000). The ubiquitination state of the ribosome provides a clue to understand translational regulation in several biological processes. However, little is known about the dynamics of ribosome ubiquitination during the life cycle of multicellular animals.

Biochemical analysis of ribosomes is often challenging in multicellular organisms, as samples from multiple tissues or developmental time points need to be processed in parallel. Ribosomes are often purified using ultracentrifugation in a gradient sucrose (Johannes et al. 1999; Reschke et al. 2013). Although effective, this method necessitates long experimental durations and substantial quantities of biological materials, and changing the purifying conditions is challenging. An alternate method uses affinity purifying ribosomes with a tagged ribosomal protein that expressed either exogenously or endogenously (Inada et al. 2002; Sanz et al. 2009; Shi et al. 2017; Simsek et al. 2017; Tryon et al. 2013). This approach enables the purification of fully assembled 80S ribosomes in a relatively short duration and with a small amount of sample. In order to understand the complicated state of ribosome modifications in the life cycle of multicellular organisms, it will be beneficial to develop an affinity purification system of the ribosome in model animals.

Here, we used zebrafish as a model system to analyze the dynamics of ribosome ubiquitination during development. We generated a zebrafish strain expressing a FLAG-tagged ribosomal protein Rpl36/eL36 from the endogenous locus. We purified the ribosomes from zebrafish embryos and larvae using this strain, and we examined the temporal change of ribosome ubiquitination level during zebrafish developed. Present study reveals that Rps10/eS10 ubiquitination by Znf598 contributes to establishing a characteristic ubiquitination pattern during zebrafish development.

## Results

### Establishment of a ribosome affinity purification system in zebrafish

To purify ribosomes from zebrafish embryos, we set up an affinity purification system with FLAG-tagged Rpl36/eL36 reported originally in mammalian cells (Simsek et al. 2017). We first verified the utility of this strategy in zebrafish embryos by injecting an mRNA encoding C-terminally FLAG-tagged Rpl36/eL36 (Rpl36-FLAG). As expected, transiently expressed Rpl36-FLAG was incorporated into 80S ribosomes, enabling us to purify both the 40S and 60S subunits by FLAG-immunoprecipitation (Figure S1A–C). Then, using CRISPR-Cas9-mediated genome editing, we inserted a FLAG-tag sequence to the 3’ end of the endogenous zebrafish *rpl36* ORF in order to stabilize the expression of Rpl36-FLAG. As a result, we obtained a strain carrying an edited *rpl36* gene locus with FLAG-tag sequence (Figure 1A). The *rpl36*-FLAG allele was transmitted to founders at the expected Mendelian ratio, suggesting that the inserted FLAG-tag sequence did not affect viability. Unless noticed, we used embryos obtained by crossing *rpl36*-FLAG heterozygous fish in this study (hereafter called *rpl36*-FLAG embryos). Fractionation of *rpl36*-FLAG embryonic lysate showed that Rpl36-FLAG was incorporated in the functioning 60S subunit: Rpl36-FLAG was detected in 60S, 80S monosome, and polysome fractions (Figure 1B). After that, we carried out FLAG-immunoprecipitation using 24-hours post fertilization (hpf) embryos, and we verified the purification of ribosomes using three methods. First, electrophoresis of the immunoprecipitated RNA revealed that both 18S and 28S rRNAs were purified from *rpl36*-FLAG embryos (Figure 1C). Second, immunoblotting analysis showed that a large ribosomal subunit protein Rpl7a/eL8 and a small ribosomal subunit protein Rps10/eS10 were copurified with Rpl36-FLAG (Figure 1D). Third, protein staining showed that proteins that were less than 50 kDa, which is the size of the most of ribosomal proteins (Martini and Gould 1975) (Figure 1E) were copurified with Rpl36-FLAG. Hence, the established *rpl36*-FLAG strain enabled us to purify ribosomes from zebrafish embryos by FLAG-immunoprecipitation.

**Figure 1.**
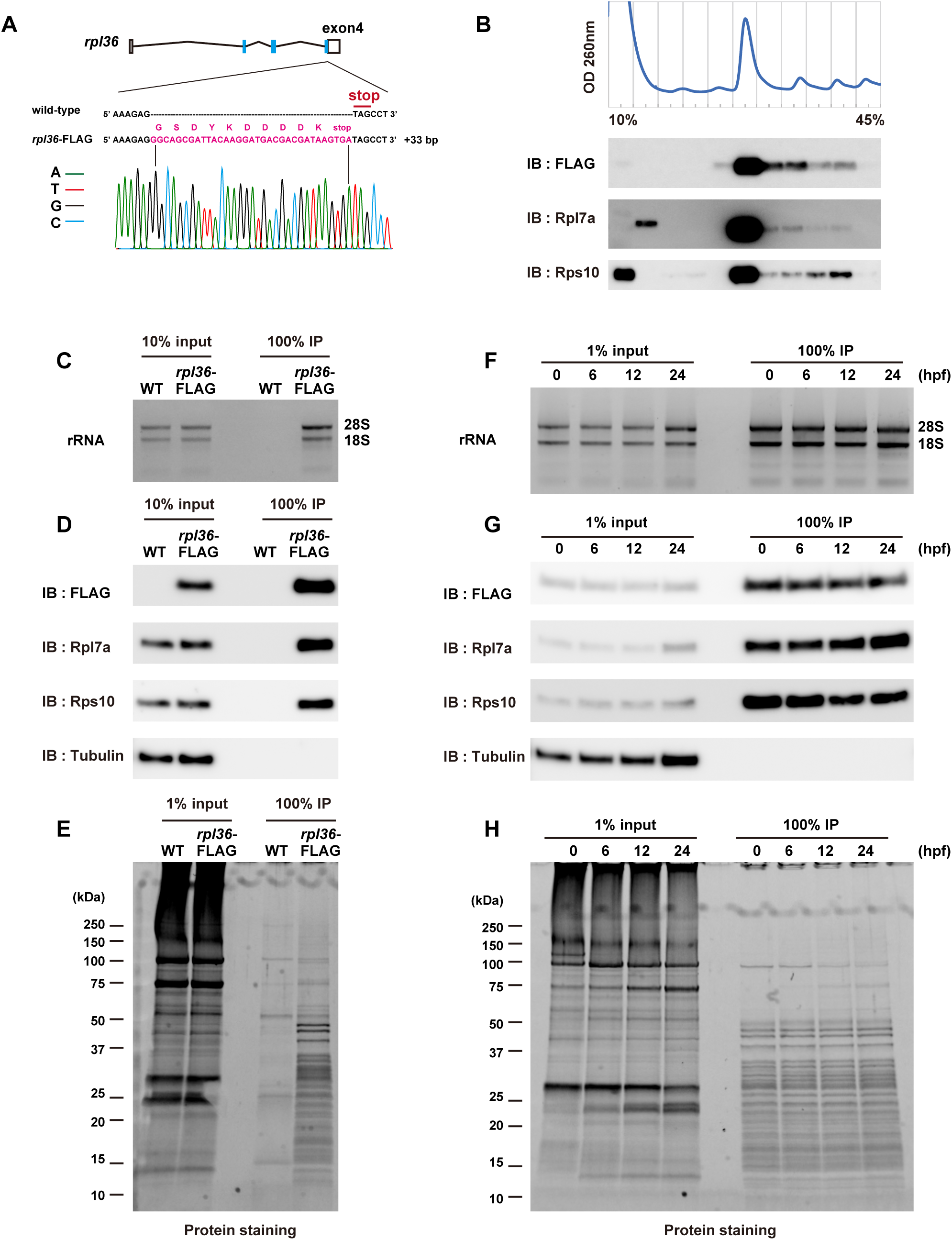
Establishment of a ribosome affinity purification system in zebrafish. (A) Scheme of a FLAG-tagged *rpl36* gene locus. Genome sequences around the stop codon of wild-type and *rpl36*-FLAG strains are shown. The sequence chromatogram of *Rpl36*- FLAG embryos is indicated below. (B) Representative polysome profiles of *rpl36*-FLAG embryos at 24 hpf. The distribution pattern of ribosomal proteins are indicated below. (C-E) Validation of the ribosome purification system with wild-type (WT) and *rpl36*-FLAG embryos at 24 hpf. Total lysates (input) and FLAG-immunoprecipitants (IP) were subjected to RNA electrophoresis (C), immunoblotting analysis using antibodies against the indicated proteins (D), and protein staining (E). (F-H) Validation of the ribosome purification system during various developmental stages. Developmental stage (in hours post-fertilization (hpf)) is indicated above. Total lysates (input) and FLAG-immunoprecipitants (IP) were subjected to RNA electrophoresis (F), immunoblotting analysis using antibodies against the indicated proteins (G), and protein staining (H).

We carried out FLAG-immunoprecipitation using the same number of embryos at 0, 6, 12, and 24 hpf to verify if our ribosome purification system is applicable to samples from various developmental stages. Ribosomal components were detected in the FLAG- immunoprecipitants at comparable levels throughout all developmental stages (Figure 1F– H). Mass spectrometry analysis of the FLAG-immunoprecipitants detected the most ribosomal proteins of small and large subunits in the all developmental stages (Figure S2A). This analysis also detected various non-ribosomal proteins involved in translation, such as Pabpc1a, Ddx6, and eEF1a (Figure S2B and Table S1). Immunoblotting analysis further confirmed copurification of these proteins with the FLAG-immunoprecipitated ribosomes (Figure S2C). These findings suggest that we can purify the ribosome complex engaging in translation from zebrafish embryos at various developmental stages by FLAG-immunoprecipitation using the *rpl36*-FLAG strain.

### Detection of ribosome ubiquitination under normal and stress conditions

After developing a zebrafish ribosome purification system, we considered this system to analyze the ubiquitination status of the ribosome. We performed immunoblotting analysis using an anti-Ubiquitin antibody on FLAG-immunoprecipitants obtained from 24 hpf wild-type and *rpl36*-FLAG embryos. Several ubiquitination signals were detected in the FLAG-immunoprecipitant from *rpl36-*FLAG embryos (Figure 2A). As mentioned above, the FLAG-immunoprecipitant contains not only ribosomes but also ribosome- or mRNA-associated proteins. Therefore, we performed FLAG-immunoprecipitation using the high-salt wash buffer to minimize the copurification of ribosome- and mRNA-associated factors (Figure S3A). When FLAG-immunoprecipitant was washed vigorously, we found that Ddx6 and eEF1a were not copurified, but the ubiquitination signals remained the same (Figure S3A and B). Despite the possibility that some ubiquitination signals originate from elements that are closely associated with the ribosome, as shown by Pabpc1 (Figure S3A), we assumed that the majority of signals were caused by the ubiquitination of core ribosomal proteins.

**Figure 2.**
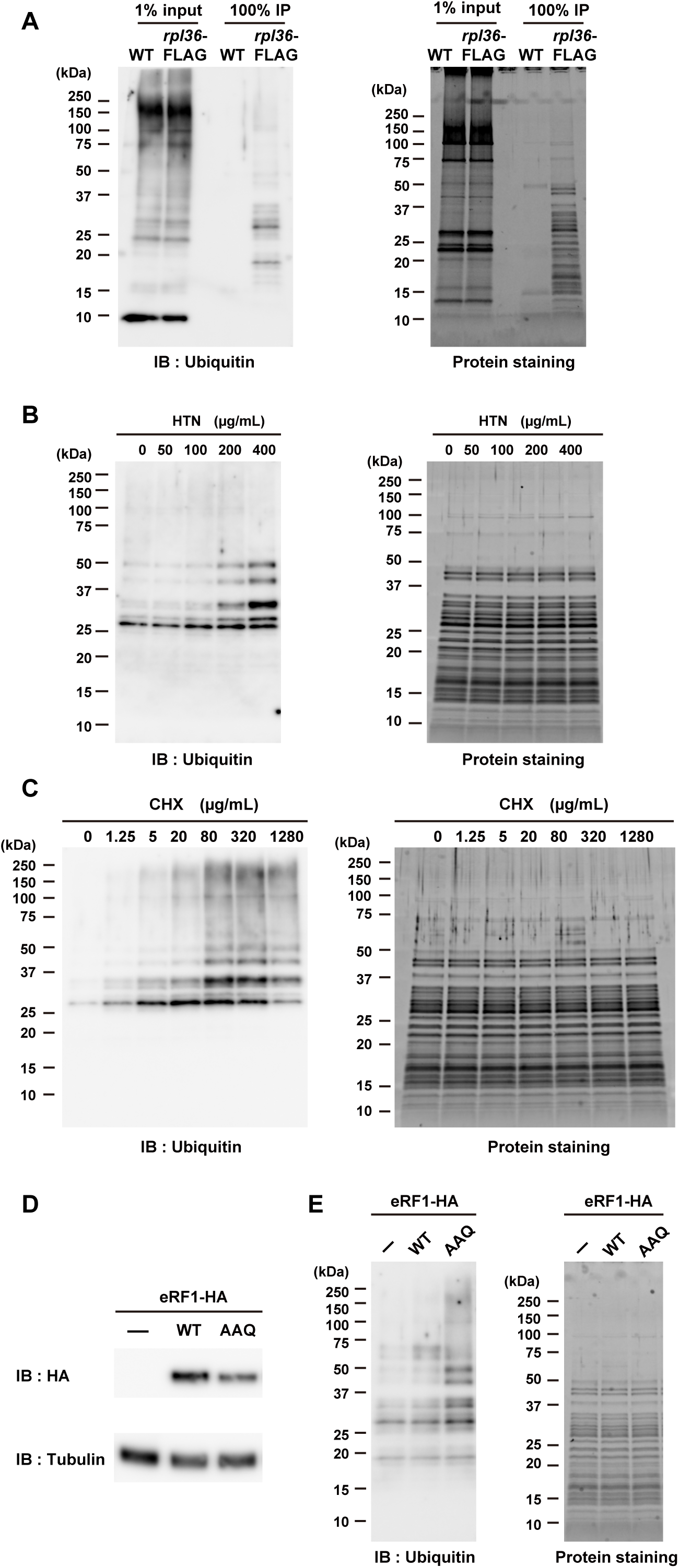
Detection of ribosome ubiquitination under normal and stress conditions. (A) Detection of ribosome ubiquitination. Total lysates (input) and FLAG- immunoprecipitants (IP) obtained from wild-type (WT) and *rpl36*-FLAG embryos at 24 hpf were subjected to immunoblotting analysis with an anti-Ubiquitin antibody (left) and protein staining (right). (B) Analysis of ribosome ubiquitination under harringtonine (HTN) treatment. FLAG-immunoprecipitants were subjected to immunoblotting analysis with an anti-Ubiquitin antibody (left) and protein staining (right). HTN concentrations are indicated above. (C) Analysis of ribosome ubiquitination under cycloheximide (CHX) treatment. FLAG-immunoprecipitants were subjected to immunoblotting analysis with an anti-Ubiquitin antibody (left) and protein staining (right). CHX concentrations are indicated above. (D-E) Analysis of ribosome ubiquitination in the presence of mutant release factor eRF1-AAQ. Total lysates were subjected to immunoblotting analysis of eRF1-HA and Tubulin (D). FLAG-immunoprecipitants were subjected to immunoblotting analysis with an anti-Ubiquitin antibody (E, left) and protein staining (E, right).

Ribosomes are subjected to ubiquitination upon translational stresses (Garshott et al. 2021; Garzia et al. 2021; Higgins et al. 2015; Juszkiewicz et al. 2018; Oltion et al. 2023; Simms et al. 2017). We thus investigated whether our ribosome purification system can detect ribosome ubiquitination triggered by translation inhibitors, which are known to enhance ribosome ubiquitination. Harringtonine (HTN) causes Rps2/uS5 and Rps3/uS3 ubiquitination by inducing 80S ribosome stalling on the initiation codon (Garshott et al. 2021; Garzia et al. 2021). When we treated *rpl36*-FLAG embryos with different concentrations of HTN and analyzed FLAG-immunoprecipitants with the anti-Ubiquitin antibody, we observed a significant dose-dependent increase in ubiquitination signals (Figure 2B). We next treated *rpl36*-FLAG embryos with a translation elongation inhibitor cycloheximide (CHX). Treatment of elongation inhibitors with intermediate concentrations causes ribosome collisions and induces Rps10/eS10 and Rps3/uS3 ubiquitination (Juszkiewicz et al. 2018; Simms et al. 2017). In fact, we observed that ubiquitination signals were upregulated as CHX concentration increased, reaching the maximum intensity at 80-320 μg/mL (Figure 2C). The signals decreased at the highest concentration (1280 μg/mL), probably because the majority of ribosomes stalled at this concentration and fewer collisions were induced, as observed in other species (Juszkiewicz et al. 2018).

As an alternative approach, we took advantage of a GGQ motif mutant of eukaryotic release factor (eRF1-AAQ) that stalls the ribosome on termination codons and induces Rps10/eS10 ubiquitination (Juszkiewicz et al. 2018). Fertilized eggs from the *rpl36*-FLAG strain were injected with mRNAs encoding C-terminally Hemagglutinin (HA)-tagged wild-type Etf1b (a zebrafish orthologue of eRF1) or Etf1b-AAQ mutant (Figure 2D). As expected, ubiquitination signals increased at 6hpf when eRF1-AAQ was injected (Figure 2E). These results indicate that, like other species, zebrafish embryos exhibit ribosome ubiquitination level fluctuations in response to translational stresses. Our FLAG-immunoprecipitation system is thus suitable for monitoring the dynamics of ribosome ubiquitination occurring in zebrafish embryos.

### Ribosome ubiquitination level changes during zebrafish development

Next, we purified ribosomes from embryonic (0 hpf to 2-days post fertilization (dpf)) and larval stages (3 dpf to 7 dpf), and analyzed the dynamics of ribosome ubiquitination during zebrafish development. Ribosome ubiquitination signals temporally changed during development (Figure 3A and B). Although it was most noticeable in the signal above 25 kDa (Figure 3A, arrowhead), this tendency was present in all of the signals detected between 25 kDa and 50 kDa (Figure 3A, bracket). Immediately following fertilization (0 hpf), the level of ribosome ubiquitination was low; but as development proceed on, it gradually increased. The ubiquitination level reached the maximum intensity at 24 hpf and then decreased toward 7 dpf (Figure 3B). These findings indicate that during development, the level of ribosome ubiquitination fluctuates.

**Figure 3.**
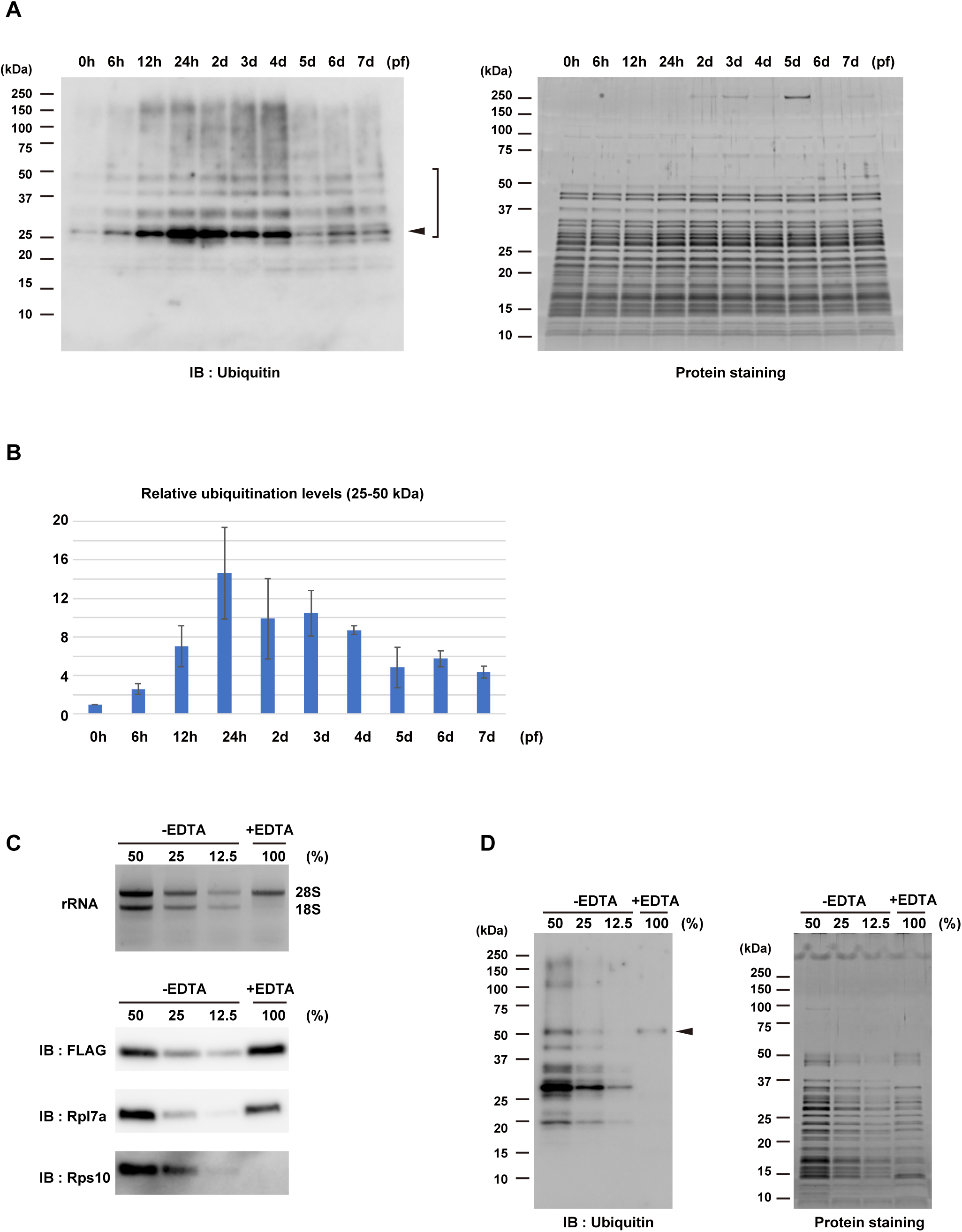
Ribosome ubiquitination level changes during zebrafish development. (A) Detection of ribosome ubiquitination during development. FLAG-immunoprecipitants from various developmental stages were subjected to immunoblotting analysis with an anti-Ubiquitin antibody (left) and protein staining (right). Arrowhead indicates most noticeable ubiquitination signal. Ubiquitination signals between 25 kDa and 50 kDa were reproducibly detected (bracket). Developmental stage is indicated above in terms of hours post-fertilization (hpf) or days post-fertilization (dpf). (B) A bar graph shows relative ubiquitination levels relative to 0 hpf. Ubiquitination signals between 25 kDa and 50 kDa in (A, left) were normalized by corresponding protein amounts in (A, right). The average of three independent experiments is indicated. (C) Validation of 60S subunits purification. FLAG-immunoprecipitants in the presence (+) or absence (-) of EDTA were subjected to RNA electrophoresis (upper) and immunoblotting analysis of ribosomal proteins (lower). The -EDTA samples were serially diluted as indicate above. (D) Detection of ribosome ubiquitination in the presence (+) or absence (-) of EDTA. FLAG-immunoprecipitants in (C) were subjected to immunoblotting analysis with an anti-Ubiquitin antibody (left) and protein staining (right). Arrowhead indicates a ubiquitination signal derived from 60S subunits.

To understand the characteristics of the ubiquitination signals, we investigated which subunit of the ribosome was linked with the detected ubiquitination signals. We performed FLAG-immunoprecipitation in the presence of EDTA to dissociate 80S ribosomes into 40S and 60S subunits. As the FLAG-tag was attached to the 60S ribosomal protein Rpl36/eL36, only 60S subunit components were purified in the presence of EDTA (Figure 3C). The amount of the purified 60S subunit reduced to approximately 25%–50% compared to the yield in the absence of EDTA, suggesting that the ribosomes purified in the absence of EDTA contained 80S ribosomes forming polysomes. We therefore compared the ubiquitination signals linked with the purified 60S (+EDTA) to that of serial dilutions of 80S (-EDTA) and found that most of the signals disappeared in the purified 60S (Figure 3D). Only the signal around 50 kDa remained in the +EDTA condition, indicating that this signal was derived from the 60S subunit (Figure 3D, arrowhead). Hence, most ubiquitination signals were associated with the 80S ribosome in a 40S subunit-dependent manner. These findings suggest that during zebrafish development, the 40S ribosomal subunit components are differently ubiquitinated.

### Znf598 promotes ribosome ubiquitination during development

Several E3 ubiquitin ligases ubiquitinate ribosomal proteins in the 40S subunit (Garzia et al. 2017, 2021; Juszkiewicz et al. 2018; Li et al. 2022; Oltion et al. 2023; Panasenko and Collart 2012; Sugiyama et al. 2019). Since we had already generated a *znf598* mutant zebrafish strain (Mishima et al. 2022), we utilized this strain to investigate whether Znf598 affects the ribosome ubiquitination during zebrafish development. We crossed the *rpl36*-FLAG strain with the *znf598* mutant strain and obtained fish carrying heterozygous *rpl36*-FLAG allele and homozygous *znf598* mutant allele. Crossing female and male of this genotype, we obtained maternal-zygotic *znf598* (MZ*znf598*) mutant embryos expressing Rpl36-FLAG (hereafter called MZ*znf598*; *rpl36*-FLAG embryos). The ubiquitination status of ribosomes purified from MZznf598; *rpl36*-FLAG embryos and *rpl36*-FLAG embryos at 0 and 24 hpf was examined. Although the initial ubiquitination levels at 0 hpf were comparable, the increase of the ubiquitination level at 24 hpf was attenuated in MZ*znf598*; *rpl36*-FLAG embryos (Figure 4A and B). On the other hand, we investigated whether overexpressing Znf598 increased the ubiquitination level of ribosomes. Ubiquitination level was increased by overexpression of wild-type Znf598 but not by RING domain-mutated Znf598 (cysteines 13 and 16 were substituted with alanines; C13/16A) (Figure 4C and D). These results indicate that Znf598 enhances ribosome ubiquitination during zebrafish development.

**Figure 4.**
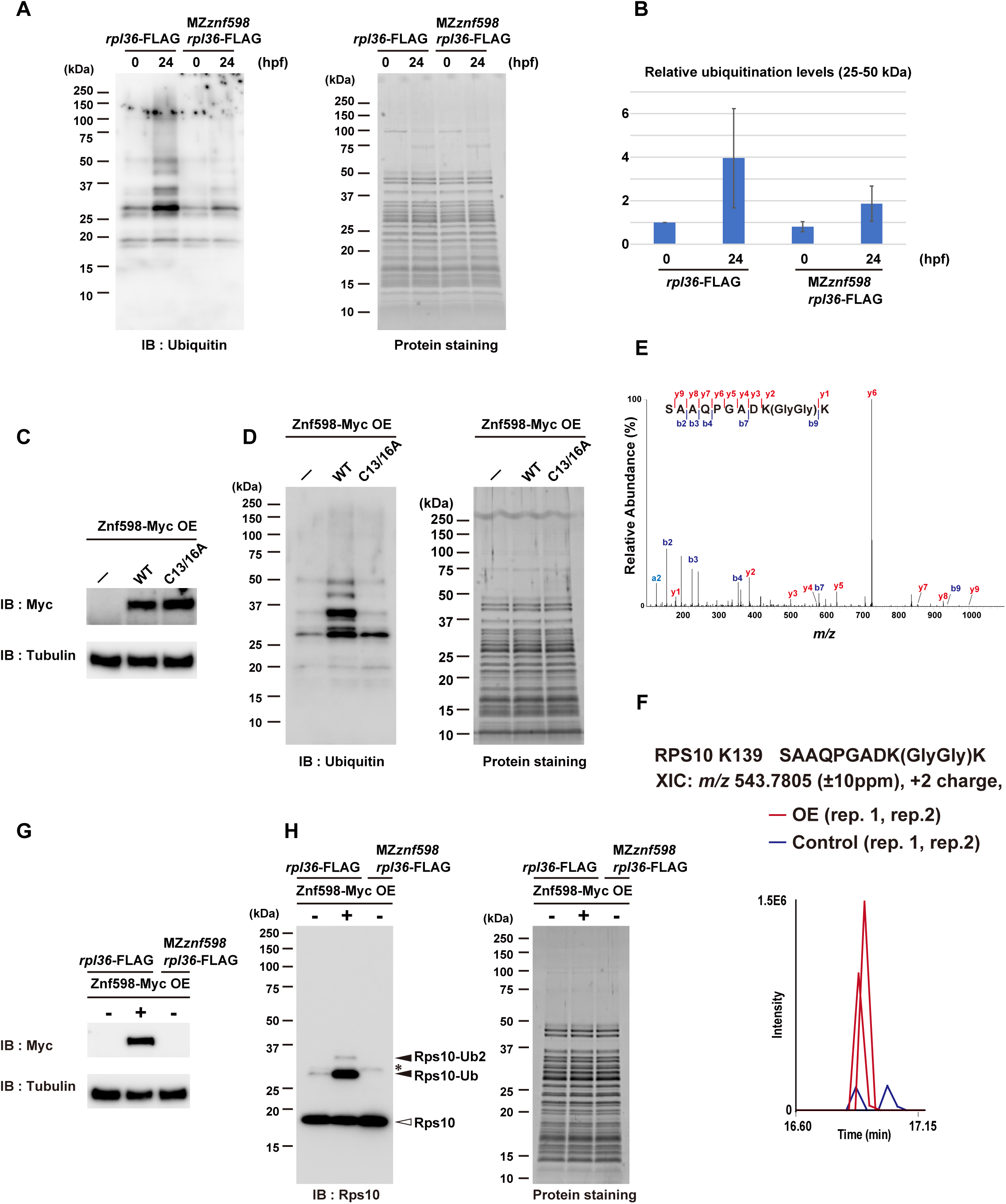
Znf598 promotes ribosome ubiquitination during development. (A) Comparison of ribosome ubiquitination levels of *rpl36*-FLAG and MZ*znf598*; *rpl36*- FLAG embryos. FLAG-immunoprecipitants from 0 and 24 hpf embryos were subjected to immunoblotting analysis with an anti-Ubiquitin antibody (left) and protein staining (right). Developmental stage (in hours post-fertilization (hpf)) is indicated above. (B) A bar graph shows ubiquitination levels relative to that of *rpl36*-FLAG embryos at 0 hpf. Ubiquitination signals between 25 kDa and 50 kDa in (A, left) were normalized by corresponding protein amounts in (A, right). The average of three independent experiments is indicated. (C-D) Comparison of ribosome ubiquitination levels with or without Znf598 overexpression. Total lysates were subjected to immunoblotting analysis of Znf598-Myc and Tubulin (C). FLAG-immunoprecipitants were subjected to immunoblotting analysis with an anti- Ubiquitin antibody (D, left) and protein staining (D, right). (E) A representative MS/MS spectrum of Rps10/eS10 di-glycyl K139. (F) MS-based quantification of a ubiquitinated peptide of Rps10/eS10 containing di-glycyl K139 under Znf598 OE (red) and control (blue) conditions. (G-H) Detection of Rps10/eS10 ubiquitination. Total lysates were subjected to immunoblotting analysis of Znf598-Myc and Tubulin (G). FLAG-immunoprecipitants were subjected to immunoblotting analysis with an anti-Rps10 antibody (H, left) and protein staining (H, right). White and black arrowheads indicate non-ubiquitinated or ubiquitinated Rps10/eS10, respectively. The asterisk indicates a non-specific signal.

### Znf598 promotes Rps10/eS10 ubiquitination in zebrafish embryos

Next, we tried identifying ribosomal protein(s) that was ubiquitinated by Znf598 during zebrafish development. Due to the low ubiquitination levels of ribosomal proteins, our first mass spectrometry analysis with FLAG-immunoprecipitants obtained from 0 to 24 hpf embryos was unable to reliably identify ubiquitinated ribosomal proteins (data not shown). Therefore, we overexpressed Znf598 to maximize the ubiquitination level at 24 hpf and subjected the FLAG-immunoprecipitants to mass spectrometry analysis. We found that an Rps10/eS10 peptide with a di-glycine remnant at lysine 139 was increased in two independent Znf598 overexpression samples (Figure 4E and F). To examine whether Rps10/eS10 was indeed ubiquitinated at 24 hpf in a Znf598-dependent manner, we purified ribosomes and detected Rps10/eS10 by immunoblotting analysis in three conditions: *rpl36*- FLAG, *rpl36*-FLAG with Znf598 overexpression, and MZ*znf598*; *rpl36*-FLAG (Figure 4G and H). The major signal appeared below 20 kDa (Figure 4H), consistent with the predicted molecular weight of zebrafish Rps10/eS10 (18.9 kDa). Additionally, in *rpl36*- FLAG embryos, a minor signal above 25 kDa was detected, which we ascribed to monoubiquitinated Rps10/eS10. Overexpression of Znf598 increased this minor signal and induced an additional slower migrating signal attributable to Rps10/eS10 attached with two ubiquitin molecules. By contrast, these additional signals were not detected in MZ*znf598*; *rpl36*-FLAG embryos. The same result was observed with exogenously expressed Rps10/eS10 with an HA-tag at the C-terminus (Figure S4A and B). These results indicate that Rps10/eS10 is ubiquitinated at 24 hpf in a Znf598-dependent manner.

While our mass spectrometry analysis detected lysine 139 in Rps10/eS10 as a potential ubiquitination site by Znf598 in zebrafish, lysines 138 and 139 (corresponding to lysines 139 and 140 in zebrafish) are ubiquitinated in a Znf598-dependent manner in mammals (Garzia et al. 2017; Sundaramoorthy et al. 2017). To validate ubiquitination site(s) in zebrafish Rps10/eS10 further, we substituted lysine 139 and/or 140 with arginine in Rps10- HA and overexpressed them in *rpl36*-FLAG embryos. We found that the ubiquitinated signal of Rps10/eS10 was not detectable in K139 single and K139/140R double mutants (Figure S4C). In contrast, the K140R single mutant reduced the ubiquitinated signal only marginally. We conclude that lysine 139 in Rps10/eS10 is a major ubiquitination site by Znf598 in zebrafish embryos, although we do not rule out the possibility that lysine 140 serves as an additional ubiquitination site.

### During zebrafish development, Rps10/eS10 ubiquitination is crucial for establishing the ribosome ubiquitination pattern

To characterize the contribution of Rps10/eS10 ubiquitination to the overall ribosome ubiquitination pattern during development, we substituted lysine 139 and 140 codons of the endogenous *rps10* locus with arginine codons by CRISPR-Cas9-mediated genome editing (*rps10* K139/140R) (Figure 5A). Homozygous *rps10* K139/140R fish reached adulthood, and we obtained a homozygous *rps10* K139/140R strain with a heterozygous *rpl36*-FLAG allele. Consistent with the experiments using Rps10-HA (Figure S4C), the ubiquitinated signals of endogenous Rps10/eS10 were not detected in ribosomes purified from the *rps10* K139/140R; *rpl36*-FLAG embryos even with Znf598 overexpression (Figure 5B and C). We then compared the overall ubiquitination pattern of purified ribosomes at 24 hpf. Similar to the ribosomes purified from MZ*znf598*; *rpl36*- FLAG embryos, multiple ubiquitination signals were reduced in ribosomes purified from *rps10* K139/140R embryos compared to ribosomes containing wild-type Rps10/eS10 (Figure 5D). Considering the molecular weight of the Rps10/eS10 ubiquitinated forms detected in the Rps10/eS10 immunoblotting analysis (Figure 5C), two ubiquitination signals were attributable to ubiquitinated forms of Rps10/eS10 (Figure 5D, black arrowheads). Indeed, the signal probably derived from the Rps10/eS10 with two ubiquitin molecules was not detected in *rps10* K139/140R; *rpl36*-FLAG and MZ*znf598*; *rpl36*- FLAG embryos. The signal probably derived from the Rps10/eS10 with monoubiquitination was reduced but still present in both *rps10* K139/140R; *rpl36*-FLAG and MZ*znf598*; *rpl36*-FLAG, likely due to an overlap with other ubiquitinated protein(s). Notably, intensities of three additional signals were also reduced in *rps10* K139/140R; *rpl36*-FLAG and MZ*znf598*; *rpl36*-FLAG (Figure 5D, white arrowheads). Although we were unable to reveal the identity of these signals, it is plausible that there are additional ribosomal proteins whose ubiquitination depends on Znf598-mediated Rps10/eS10 ubiquitination (see discussion). To further test this possibility, we compared ubiquitination signals in *rpl36*-FLAG and *rps10* K139/140R; *rpl36*-FLAG embryos under the Znf598- overexpressed condition (Figure S5A). Whereas multiple ubiquitination signals were enhanced by overexpression of Znf598 in *rpl36*-FLAG embryos, increase of these signals was significantly suppressed in *rps10* K139/140R; *rpl36*-FLAG embryos. This result further supports our assumption that Rps10/eS10 ubiquitination is a prerequisite for multiple ubiquitination events promoted by Znf598.

**Figure 5.**
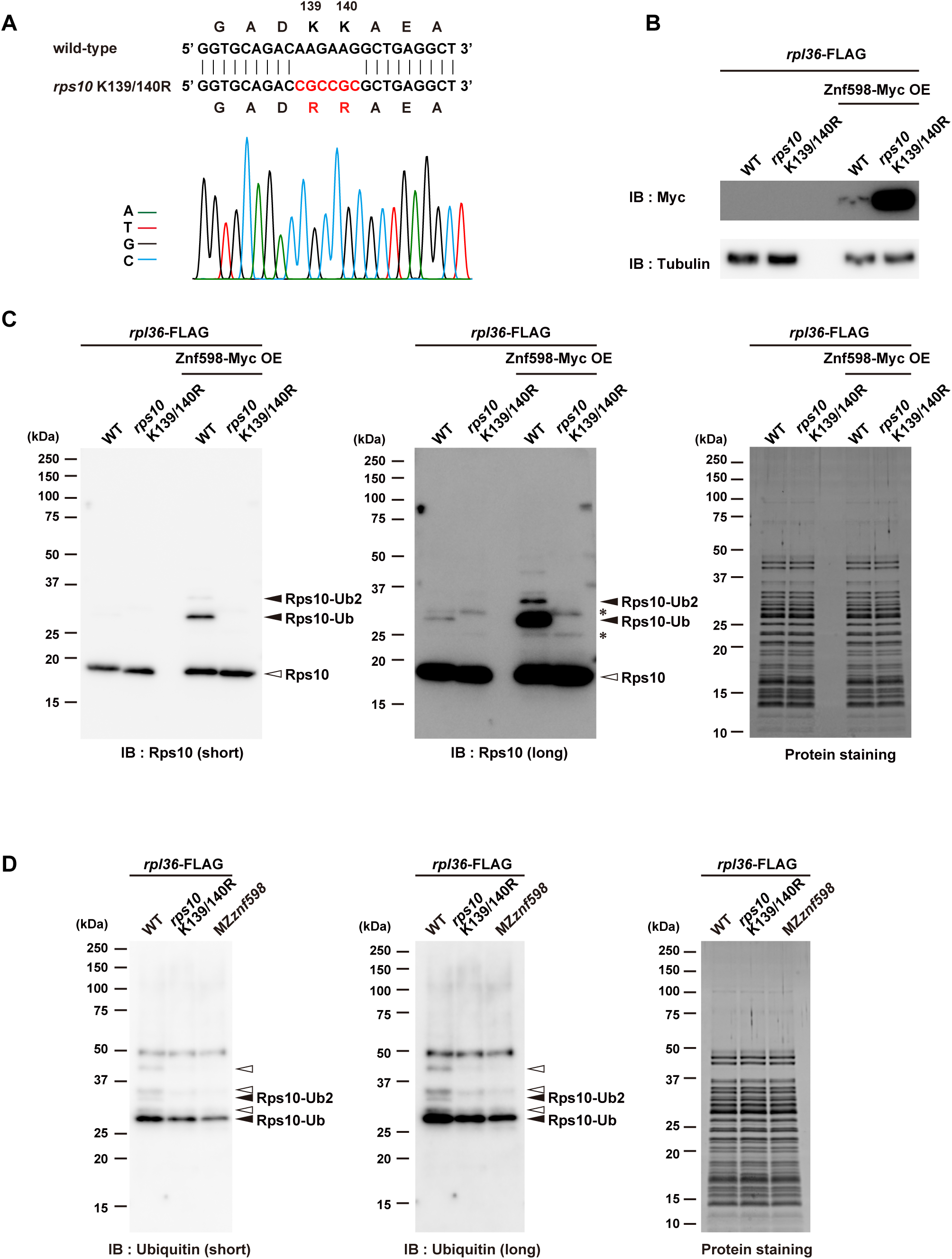
During zebrafish development, Rps10/eS10 ubiquitination is crucial for establishing the ribosome ubiquitination pattern. (A) Sequence alignment of an *rps10* gene obtained from wild-type and the *Rps10* K139/140R strain. The sequencing chromatogram of the *Rps10* K139/140R embryos is indicated below. (B-C) Detection of Rps10/eS10 ubiquitination with or without Znf598 overexpression in *rpl36*-FLAG and *rps10* K139/140R; *rpl36*-FLAG embryos. Total lysates were subjected to immunoblotting analysis of Znf598-Myc and Tubulin (B). FLAG- immunoprecipitants were subjected to immunoblotting analysis with an anti-Rps10 antibody (C, left and middle) and protein staining (C, right). White and black arrowheads indicate non-ubiquitinated or ubiquitinated Rps10/eS10, respectively. Asterisks indicate non-specific signals. (D) Comparison of ribosome ubiquitination pattern in *rpl36*-FLAG, *rps10* K139/140R; *rpl36*-FLAG, and MZ*znf598*; *rpl36*-FLAG embryos. FLAG- immunoprecipitants were subjected to immunoblotting analysis with an anti-Ubiquitin antibody (left and middle) and protein staining (right). Black arrowheads indicate putative Rps10/eS10 ubiquitinated signals. White arrowheads indicate reduced ubiquitinated signals in *rps10* K139/140R; *rpl36*-FLAG and MZ*znf598*; *rpl36*-FLAG embryos.

Given the critical role of Rps10/eS10 ubiquitination by Znf598 for establishing ribosome ubiquitination patterns at 24 hpf, we wondered if Rps10/eS10 ubiquitination *per se* fluctuates during development. To test this possibility, we performed immunoblotting analysis to examine Rps10/eS10 ubiquitination from 0 to 24 hpf. At 0 hpf, the monoubiquitinated Rps10/eS10 signal was barely detectable; however, it gradually increased until 24 hpf in rpl36-FLAG embryos, mirroring the increase in the whole ribosome ubiquitination levels (Figure 6A and Figure 3A). These Rps10/eS10 monoubiquitinated signals were completely abolished in *rps10* K139/140R; *rpl36*-FLAG embryos. Next, we analyzed Rps10/eS10 ubiquitination in larval stages. The Rps10/eS10 monoubiquitination levels peaked between 1 and 3 dpf and gradually decreased toward 7 dpf (Figure 6B), similar to the whole ribosome ubiquitination levels (Figure 3A). Overall, these results demonstrate that Rps10/eS10 ubiquitination by Znf598 occurs dynamically after fertilization and contributes directly and indirectly to establish the overall ribosome ubiquitination pattern during zebrafish development.

**Figure 6.**
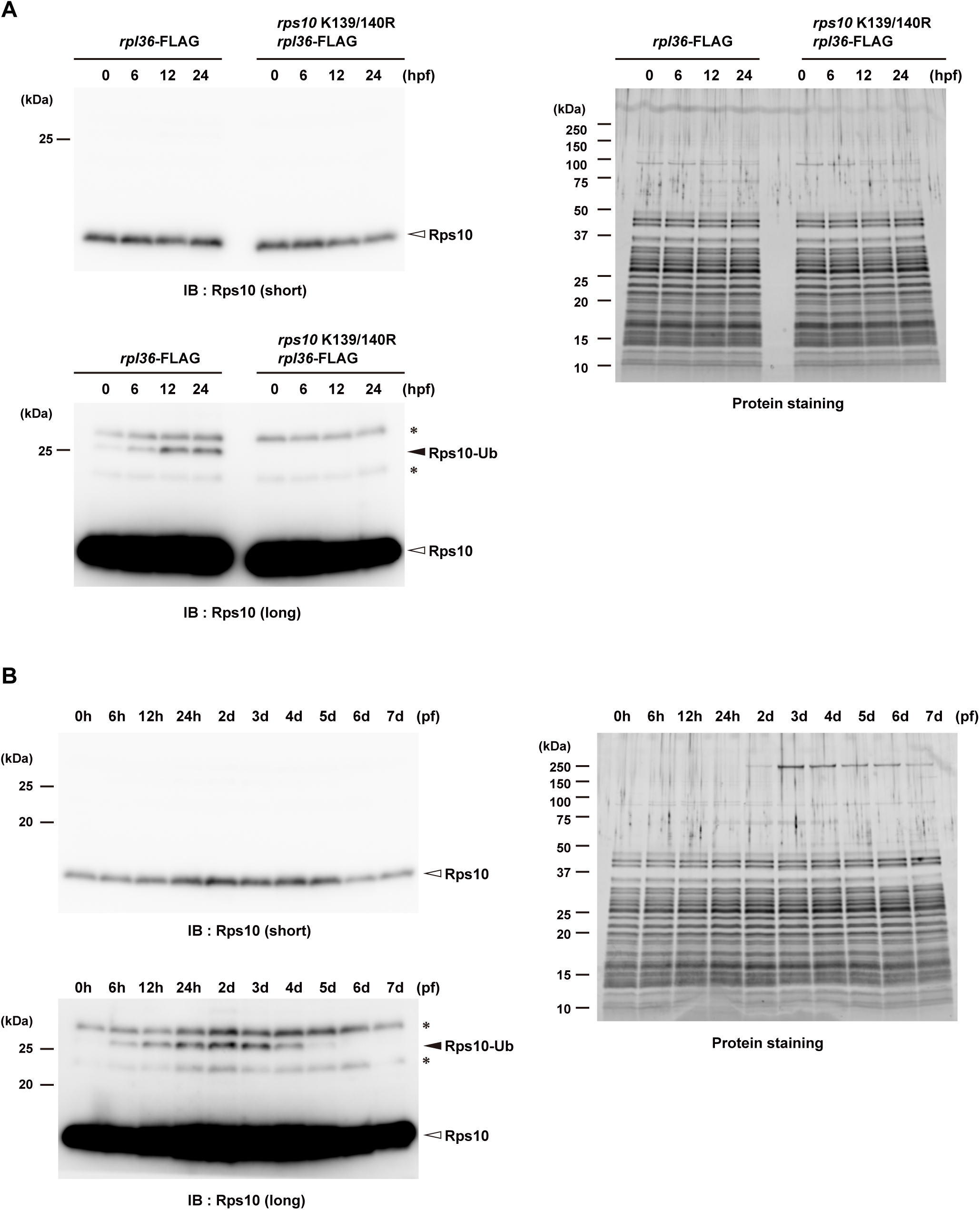
Rps10/eS10 ubiquitination level temporally changes during development. (A) Detection of Rps10/eS10 ubiquitination from 0 to 24 hpf embryos. FLAG- immunoprecipitants were subjected to immunoblotting analysis with an anti-Rps10 antibody (left) and protein staining (right). Developmental stage (in hours post-fertilization (hpf)) is indicated above. White and black arrowheads indicate non-ubiquitinated or ubiquitinated Rps10/eS10, respectively. Asterisks indicate non-specific signals. (B) Detection of Rps10/eS10 ubiquitination during development. FLAG-immunoprecipitants were subjected to immunoblotting analysis with an anti-Rps10 antibody (left) and protein staining (right). Developmental stage (in hours post-fertilization (hpf) or days post- fertilization (dpf)) is indicated above. White and black arrowheads indicate non- ubiquitinated or ubiquitinated Rps10/eS10, respectively. Asterisks indicate non-specific signals.

## Discussion

In this study, we established a novel ribosome affinity purification system in zebrafish by introducing a FLAG-tag into endogenous Rpl36/eL36. Using this system, we affinity purified almost fully assembled 80S ribosomes from zebrafish embryos at different developmental stages. In comparison to the standard ribosome purification methods employing a gradient sucrose and ultracentrifugation, this method offers certain advantages due to the simplicity of the experimental process, good repeatability, and efficiency (20% of the input).

Transgenic zebrafish strains expressing an exogenous copy of EGFP-fused Rpl10a/uL1 were generated previously and were utilized for purifying ribosome-associated mRNAs (Tryon et al. 2013). However, engineering an endogenous ribosomal gene locus with an affinity tag insertion has been challenging in zebrafish. Our *rpl36*-FLAG strain is the first example of tagging an endogenous ribosomal protein in zebrafish, enabling the purification of ribosomes with minimum perturbation of the stoichiometry of ribosomal proteins. Rpl36-FLAG was incorporated into the 60S subunit efficiently and allowed the purification of 80S ribosomes to analyze ubiquitination status. Our *rpl36*-FLAG strain will assist in the analysis of ribosome modifications as well as ribosome components, ribosome-associated proteins, and ribosome-associated mRNAs during zebrafish development, even though it is uncertain whether the ribosome containing Rpl36-FLAG is functionally identical to the wild-type ribosome.

Combining the ribosome affinity purification approach with genetic mutants, we revealed a complex and dynamic nature of ribosome ubiquitination during zebrafish development. The ubiquitination level of ribosomes was at its lowest just after fertilization, indicating that ribosomes in unfertilized eggs are mostly stored without ubiquitination. Before fertilization, global translation is silenced, and ribosomes are kept in a dormant state (Leesch et al. 2023). The cotranslational ubiquitination pathways should not be at work in such a situation. Hence, the low level of ribosome ubiquitination in fertilized eggs may reflect the absence of cotranslational ribosome ubiquitination during oogenesis. Alternatively, but not exclusively, the activity of ribosome deubiquitinating enzymes such as USP10, OTUD1, USP21, and OTUD3 (Garshott et al. 2020; Meyer et al. 2020; Snaurova et al. 2022) may predominate over ribosome ubiquitination activity during oogenesis, keeping the ribosome less ubiquitinated. In any case, our data indicate that the maternal ribosome starts from a naïve ubiquitination state in zebrafish embryogenesis.

In the present study, we revealed that multiple ribosome ubiquitination signals gradually increased after fertilization and ubiquitination of Rps10/eS10 by Znf598 contributed to this process. As Znf598 plays a pivotal role in RQC and its ubiquitination sites in Rps10/eS10 are highly conserved (Garzia et al. 2017; Juszkiewicz and Hegde 2017; Sundaramoorthy et al. 2017), it is likely that the ribosome ubiquitination increases after fertilization in response to ribosome collisions. It is possible that following fertilization, translation activation induces ribosome collisions on stall-prone mRNAs, and that as collided ribosomes accumulate, Znf598-dependent ribosome ubiquitination increases. Supporting this scenario, ribosome collision sites were detected in more than 1000 mRNA species at 4 hpf (Han et al. 2020). Alternatively, Znf598 may be developmentally regulated. Compared to the lysate of HEK293, Znf598 accumulates at significantly lower levels in rabbit reticulocytes (Juszkiewicz et al. 2018), indicating that Znf598 steady-state levels may vary depending on the cellular situation. Considering the sub-stoichiometric abundance of Znf598 (Garzia et al. 2017) and Znf598 overexpression in zebrafish embryos increased ribosome ubiquitination at 24 hpf, the Znf598 protein amount can be a major determinant of ribosome ubiquitination levels in response to ribosome collisions. During oxidative stress, inhibition of a deubiquitinating enzyme promote ribosome ubiquitination (Silva et al. 2015). Therefore, the degree of deubiquitinating enzyme(s) activation or expression is also a critical factor to promote ribosome ubiquitination. Future research must determine the frequency of ribosome collision and the balance between the expression and activation of the deubiquitinating enzyme(s) and Znf598 during zebrafish development.

A balance between ZNF598-mediated ubiquitination and other molecular pathways that recognize collided ribosomes are another factor that potentially affects the ubiquitination levels of the ribosome. ZAKα is recruited to collided ribosomes and phosphorylates p38 and JNK, leading to cell death (Wu et al. 2020; Vind et al. 2020). Additionally, ZAKα recruits GCN1 to colliding ribosomes (Wu et al. 2020), and GCN1 further binds to colliding ribosomes to provide a scaffold for eIF2 phosphorylation, therefore attenuating global translation initiation and preventing further ribosome collision (Pochopien et al. 2021; Yan and Zaher 2021). EDF1 is an additional collided ribosome sensor and inhibits translation initiation via GIGYF2-4EHP complex (Juszkiewicz et al. 2020a). Therefore, competition among Znf598, ZAKα, GCN1, and EDF1 on the collided ribosomes may impact the effectiveness of ribosome ubiquitination. In addition, several factors are reported to reduce ribosome stalling, such as GTPBP2 and eIF5A (Ishimura et al. 2014; Schuller et al. 2017). We assume that during development, a delicate balance between these multiple factors is not static but rather dynamic and contributes to the changes in the ubiquitination levels of ribosome.

Although, Rps10/eS10 is one of the key ubiquitinated ribosomal proteins in zebrafish embryos, neither loss of Znf598 nor mutations of the ubiquitination sites in Rps10/eS10 abolished ubiquitination signals on the ribosome completely, indicating that Rps10/eS10 alone cannot explain the entire ubiquitination pattern of the ribosome during development. The identity of the remaining ubiquitinated protein(s) needs to be determined. Our data showed that most of the ubiquitination signals were linked with 40S subunit of ribosome. Furthermore, Znf598 mutant and Rps10 K139/140R mutant not only abolished the Rps10/eS10 ubiquitination *per se*, but also reduced the additional ubiquitination signals on the ribosome. These findings imply that further ubiquitination events on the 40S subunit require Znf598-mediated Rps10/eS10 ubiquitination as a prerequisite. Consistent with our results, ubiquitination levels of Rps20/uS10 and Rps3/uS3 are reduced in the absence of Rps10/eS10 ubiquitination in mammalian cells (Garshott et al. 2020; Meyer et al. 2020). Similarly, it has been observed that ubiquitination of Rps3/uS3 is necessary for ubiquitination of Rps2/uS5 (Meyer et al. 2020). These observations and our data collectively suggest that the 40S subunit is ubiquitinated hierarchically, and Znf598-mediated Rps10/eS10 ubiquitination plays a key role in the process. Future research should combine immunoblotting analyses with antibodies against these candidate ribosomal proteins and the generation of strains with ubiquitination-site mutations to ascertain the ubiquitination state of these ribosomal proteins during development.

One remaining question is the biological relevance of the ribosome ubiquitination dynamics to development. Both the homozygous *znf598* mutant fish and the homozygous *rps10* K139/140R fish had morphologies similar to those of wild-type fish and had the ability to create founders on their own. Hence, the requirement of Znf598-mediated Rps10/eS10 ubiquitination during zebrafish development is still unknown. Future research is necessary to identify morphologically undetectable abnormalities in the absence of Znf598-mediated Rps10/eS10 ubiquitination and to determine if the trait develops under translationally stressful conditions.

## Materials and methods

### Zebrafish husbandry

The zebrafish AB strain was used as a wild-type stain and maintained according to the animal experiment protocol (2018-46) at Kyoto Sangyo University. Fishes were raised and maintained at 28.5°C under standard laboratory conditions with the cycle of 14 hour-light and 10 hour-dark. Natural breeding method was used to produce fertilized eggs and embryos were grown in system water at 28.5°C.

### Generation of *rpl36*-FLAG and *rps10* K139/140R strains

The zebrafish *rpl36*-FLAG and *rps10* K139/140R strains were generated by CRISPR-Cas9-mediated genome editing. CRISPRscan (Vejnar et al. 2016) was used to predict a guide RNA targeting the exon 4 of the *rpl36* gene locus (ENSDARG00000100588) or exon 5 of the *rps10* gene locus (ENSDARG00000034897). The DNA template for sgRNA synthesis was prepared by PCR using the gene specific sgRNA primer and the sgRNA tail primer as shown in Table S2. The template was transcribed with T7 RNA Polymerase (TAKARA) and purified using Probe-Quant G-25 microcolumns (Cytiva). As shown in Table S2, the single-stranded oligodeoxynucleotides (ssODNs) were synthesized by Eurofins Genomics. The recombinant Cas9 protein (TAKARA) and the sgRNA were incubated for 10 min at 37°C, and co-injected with the ssODN in the one-cell zebrafish embryos (0.45 µg/µl Cas9 protein, 90 ng/µl sgRNA, 0.125 pmol/µl ssODN). Injected embryos were grown into adult fish and crossed with wild-type fish. Using the appropriate set of primers (Table S2), embryos carrying an insertion/substitution in the corresponding gene locus were screened by genotyping PCR, and siblings with potential insertion/substitution were raised. Fin-clipping was used to genotype adult fish, and Sanger sequencing was used to verify that the editing was successful.

### Generation of MZ*znf598*; *rpl36*-FLAG and *rps10* K139/140R; *rpl36*-FLAG strains

The *znf598* mutant strain generated previously (Mishima et al. 2022) and the *rps10* K139/140R mutant strain generated in this study were crossed with the *rpl36*-FLAG strain. Heterozygous fish were crossed to generate homozygous *znf598* mutant or *rps10* K139/140R mutant in the *rpl36*-FLAG heterozygous background.

### Plasmid construction

The primer sequences and plasmids used in this study are shown in Tables S2 and S3, respectively. The ORF of zebrafish *rpl36* (ENSDARG00000100588) was amplified by RT-PCR and cloned into pCS2+ via EcoRI/XhoI restriction sites using DNA Ligation Kit (TAKARA). The ORF of zebrafish *znf598* (ENSDARG00000014945) was amplified by RT-PCR and cloned into pCS2+ via XhoI/XbaI restriction sites using DNA Ligation Kit (TAKARA). To generate *znf598* point mutation in ORF (*znf598* C13/16A), wild-type ORF was amplified using point mutated primers and cloned into pCS2+ via XhoI/XbaI restriction sites using DNA Ligation Kit (TAKARA). The ORFs of zebrafish *etf1b* (ENSDARG00000043976) and *rps10* (ENSDARG00000034897) were amplified by RT-PCR and cloned into pCS2+HA using NEBuilder HiFi DNA Assembly (New England BioLabs). The pCS2+HA was generated by inserting synthetic DNA fragments with HA-tag sequence into pCS2+ via EcoRI/XbaI restriction sites. To generate *etf1b* and *rps10* point mutations in ORFs (*etf1b*-AAQ, *rps10-*K139R, *rps10-*K140R, and *rps10-*K139/140R), wild-type ORFs were amplified using point mutated primers. DNA fragments were then assembled using NEBuilder HiFi DNA Assembly (New England BioLabs).

### Polysome analysis

For polysome analysis, we followed the original protocol with some modifications (Mishima et al. 2012). Zebrafish embryos at 24 hpf were dechorionated with 1 mg/mL pronase (Sigma). Sixty dechorionated embryos were homogenized in buffer A (20 mM HEPES-KOH pH 7.5, 100 mM KCl, 10 mM MgCl_2_, 0.25% NP-40, 250 mM sucrose, 2 mM DTT, 10 µg/mL cycloheximide, 100 U/mL RNase inhibitor (Promega), Complete Protease Inhibitor EDTA-free in nuclease-free water). Lysates were incubated for 5 min on ice and centrifuged for 5 min at 12,000 rpm at 4℃. Clarified lysates were loaded onto a continuous 10%–45% *(w/v)* sucrose gradient prepared in buffer B (20 mM HEPES-KOH pH 7.5, 100 mM KCl, 10 mM MgCl_2_, 2 mM DTT, 10 µg/mL cycloheximide, 100 U/mL RNase inhibitor (TAKARA)). Gradients were centrifuged in a P40ST rotor (himac) for 180 min at 36,000 rpm at 4°C. Polysome gradients were analyzed using a gradient station (BioComp) coupled to a TriaxTM Flow Cell detector (FC-2). Each fraction was collected and precipitated by ethanol (75% final). Pellets were dissolved in Sample Buffer Solution (Fujifilm-Wako). Samples were incubated for 30 min at 92°C and proceed to SDS-PAGE followed by immunoblotting analysis.

### Immunoblotting analysis

Samples were incubated with Sample Buffer Solution (Fujifilm-Wako) for 10 min at 92°C. The SDS-PAGE and immunoblotting experiments were carried out according to standard protocols. Signals were developed using ImmunoStar LD (Fujifilm-Wako) or Luminata Forte (Millipore) and detected using Amersham Imager 680 (GE Healthcare). Antibodies used in this study are shown in Table S4.

### Immunoprecipitation

To conduct the immunoprecipitation assay, we followed the original protocol with some modifications (Simsek et al. 2017). For immunoprecipitation by anti-FLAG antibody, 15–45 zebrafish embryos were homogenized in the lysis buffer A (25 mM Tris-HCl pH 7.6, 150 mM NaCl, 15 mM MgCl_2_, 1 mM DTT, 8% glycerol, 1% triton X-100, 50 µM PR-619, 100 U/ml RNase inhibitor (TAKARA), Complete Protease Inhibitor EDTA-free in nuclease-free water). Lysates were incubated for 5 min on ice and centrifuged for 5 min at 2,000 g at 4°C. Supernatants were incubated with Anti-DYKDDDDK tag Antibody Beads (Fujifilm-Wako) for 15 min on rotation at 4°C.

Anti-DYKDDDDK tag Antibody Beads were prepared by washing with wash buffer (25 mM Tris-HCl pH 7.6, 150 mM NaCl, 15 mM MgCl_2_, 1 mM DTT, 8% glycerol, 1% triton X-100) three times. Four micro-litter of slurry beads were used per embryo.

After 15 min incubation at 4°C with rotation, beads were washed twice with buffer B (25 mM Tris-HCl pH 7.6, 150 mM NaCl, 15 mM MgCl_2_, 1 mM DTT, 1% triton X-100, 50 µM PR-619) and once with buffer C (25 mM Tris-HCl pH 7.6, 150 mM NaCl, 15 mM MgCl_2_, 1% triton X-100, 50 µM PR-619). Samples were eluted from the beads by incubating with 0.5 mg/mL FLAG peptide (MEDICAL&BIOLOGICAL LABORATORIES CO., LTD) in buffer C at 25°C for 30 min. For EDTA-treated samples, 50 mM EDTA was used in place of 15 mM MgCl_2_ in buffers A, B, C, and wash buffer.

For mass spectrometry analysis, beads were washed twice using buffer D (25 mM Tris-HCl pH 7.6, 150 mM NaCl, 15 mM MgCl_2_) after washing with buffer B. We followed the original protocol (Imami et al. 2018, 2023) with some modifications. Beads were resuspended in PTS buffer (12 mM sodium deoxycholate (SDC), 12 mM sodium N-lauroylsarcosinate (SLS) in 100 mM Tris-HCl pH 8.0) followed by incubation for 30 min at 37°C under constant agitation. After the incubation, the supernatant was reduced with 10 mM dithiothreitol (DTT) at 37°C for 30 min and alkylated with 50 mM iodoacetamide (IAA) at room temperature for 30 min in the dark room. The samples were diluted 5 times with 50 mM ammonium bicarbonate (ABC). For Figure S2A and B, proteins were digested by 0.5 µg lysyl endopeptidase (LysC) (Fujifilm-Wako) and 0.5 µg Trypsin (Promega) at room temperature overnight. For Figure 4E and F, proteins were digested by LysC, Trypsin, or Glu-C (Promega). Next day, 500 µl ethyl acetate (Fujifilm-Wako) was added to the sample and digestion was quenched by adding 0.5% trifluoroacetic acid (TFA) (final concentration). The samples were shaken for 1 min and centrifugated at 15,000 g for 2 min at room temperature. The organic phase containing SDC and SLS was discarded. The resulting peptide solution was evaporated in a SpeedVac. The residue was resuspended in 0.1% TFA and desalted with SDB-XC Stage tips (Rappsilber et al. 2007) prior to LC-MS/MS analysis.

For high-salt washes experiments, after 15 min incubation at 4°C rotation, beads were washed twice with buffer B containing 400 mM NaCl (25 mM Tris-HCl pH 7.6, 400 mM NaCl, 15 mM MgCl_2_, 1 mM DTT, 1% triton X-100, 50 µM PR-619). Afterwards, beads were washed once with buffer C containing 400 mM NaCl (25 mM Tris-HCl pH 7.6, 400 mM NaCl, 15 mM MgCl_2_, 1% triton X-100, 50 µM PR-619). Samples were then eluted off the beads using 0.5 mg/mL FLAG peptide in buffer C containing 400 mM NaCl for 30 min at 25°C. RNA was extracted from a portion of the samples, and ribosomal RNAs were quantified by agarose gel electrophoresis, to compensate for differences in the quantities of ribosomes recovered from low and high-salt conditions. The equal amounts of ribosomes estimated from the RNA analysis were subjected to immunoblotting and protein staining.

### LC-MS/MS analysis

Nano-scale reversed-phase liquid chromatography coupled with tandem mass spectrometry (nanoLC/MS/MS) was performed on an Orbitrap Exploris 480 (related to Figure S2A and B) or an Orbitrap Fusion Lumos mass spectrometer (related to Figure 4E and F) (Thermo Fisher Scientific) connected to a Thermo Ultimate 3000 RSLCnano pump equipped with a self-pulled analytical column (150 mm length × 100 μm i.d.) (Ishihama et al. 2002) packed with ReproSil-Pur C18-AQ materials (3 μm or 1.9 μm; Dr. Maisch GmbH). The mobile phases comprised (A) 0.5% acetic acid and (B) 0.5% acetic acid and 80% ACN.

For the Orbitrap Exploris 480 system (related to Figure S2A and B), peptides were separated on self-pulled needle columns (250 mm, 100 μm ID) packed with Reprosil-Pur 120 C18-AQ1.9 μm at 50°C in a column oven. The flow rate was 400 nL/min. The flow gradient was set as follows: 5% B in 5 min, 5–19% B in 55.3 min, 19–29% B in 21 min, 29–40% B in 8.7 min, and 40–99% B in 0.1 min, followed by 99% B for 4.9 min. The electrospray voltage was set to 2.2 kV in the positive mode. The mass spectrometric analysis was carried out with the FAIMS Pro interface (Thermo Fisher Scientific). The compensation voltage (CV) was set to −40, −60, and -80 and the cycle time of each CV experiment was set to 1 s. The mass range of the survey scan was from 375 to 1,500 *m/z* with a resolution of 60,000, 300% normalized automatic gain control (AGC) target and auto maximum injection time. The first mass of the MS/MS scan was set to 120 *m/z* with a resolution of 15,000, standard AGC, and auto maximum injection time. Fragmentation was performed by HCD with a normalized collision energy of 30%. The dynamic exclusion time was set to 20 s.

The Orbitrap Fusion Lumos instrument was operated in the data-dependent mode with a full scan in the Orbitrap followed by MS/MS scans using HCD (related to Figure 4E and F). Peptides were eluted from the analytical column at a flow rate of 500 nL/min, with the following gradient: 5–10% B for 5 min, 10–40% B for 60 min, 40–99% B for 5 min, and 99% for 5 min. The applied voltage for ionization was 2.4 kV. The full scans were performed with a resolution of 120,000, the standard AGC target mode, and a maximum injection time of 50 ms. The MS scan range was *m/z* 300–1,500. The CV of the FAIMS Pro interface was set to −40, −60, and −80, and the cycle time of each CV experiment was set to 1 s. The MS/MS scans were collected in the ion trap with the rapid mode, the standard AGC target mode, and a maximum injection time of 35 ms. The isolation window was set to 1.6, and the normalized HCD collision energy was 30. Dynamic exclusion was applied for 20 s.

### Processing of proteome data

All raw data files were analyzed and processed by maxquant (version 1.6.15.0 or 1.6.17.0) (Cox and Mann 2008), and the database search was performed with Andromeda (Cox et al. 2011), which is a peptide search engine integrated into the MaxQuant environment. Searches were conducted against a zebrafish UniProt database (version 2021- 3; 46,849 protein entries) spiked with common contaminants and enzyme sequences. Search parameters included two missed cleavage sites and variable modifications such as methionine oxidation; protein N-terminal acetylation; deamidation of glutamine and asparagine residues; and diglycine of lysine residue (only ubiquitination analysis related to Figure 4E and F). Cysteine carbamidomethylation was set as a fixed modification. The enzyme was set as trypsin/P (cleaves after lysine, and arginine if followed immediately by a proline), lysC (cleaves after lysine), or Glu-C (cleaves after glutamic acid). The peptide mass tolerance was 4.5 ppm, and the MS/MS tolerance was 20 ppm. The false discovery rate (FDR) was set to 1% at the peptide spectrum match level and protein level. The ‘match between runs’ function was performed. All necessary information regarding proteomic analyses, including protein and peptide lists, were deposited in a publicly accessible repository (jPOST) (Moriya et al. 2019; Okuda et al. 2017) with the dataset identifier, PXD039560.

### Microinjection

Microinjection was performed as described in (Mishima and Tomari 2016) using IM300 Microinjector (NARISHIGE). Approximately 1,000 pL of the solution was injected per embryo within 15 min after fertilization. Embryos were developed in system water at 28.5°C.

### Protein staining

Following SDS-PAGE, the gel was immersed in Oriole solution (BioRad) for 90 min while being constantly agitated. For five minutes, the gel was rinsed twice with distilled water. Amersham Imager 680 was used to detect the signals (GE Healthcare).

### RNA extraction

RNA samples were extracted from zebrafish embryos or FLAG-immunoprecipitants using TRI Reagent (Molecular Research Center). Ethachinmate (Nippon gene) was used in ethanol precipitation to recover a small quantity of RNA.

### *In vitro* mRNA transcription

mRNA was synthesized from a linearized plasmid DNA template using the SP6 Scribe Standard RNA IVT Kit and the ScriptCap m7G Capping System (CELLSCRIPT) or mMessage mMachine SP6 (Thermo Fisher Scientific), followed by purification with the RNeasy Mini Kit (QIAGEN).

### Harringtonine and cycloheximide treatment

Embryos were incubated with the drugs at the indicated concentrations and duration at 28.5℃. 23 hpf embryos were given an hour of treatment with harringtonine (HTN) (LKT Laboratories, Inc.) at concentrations of 50, 100, 200, or 400 µg/ml. 22 hpf embryos were given two hours of treatment with cycloheximide (CHX) (Fujifilm-Wako) at concentrations of 1.25, 5, 20, 80, 320, or 1280 µg/ml.

### Quantification of ubiquitination signals

Signals were quantified using the ROI manager function of the Image J software (http://imageJ.nih.gov/ij/). Background signal was measured with the empty lane and subtracted from quantified values. Relative ubiquitination levels were calculated by dividing a ubiquitin signal by a corresponding protein staining signal. Three biological replicates were used in the experiments, and averages were computed.

## Acknowledgments

The authors thank our laboratory members for discussions and technical assistance, especially Kimi Wakabayashi for fish maintenance. We thank Toshifumi Inada for technical advice on the ribosome analysis.

This work was supported by the Japan Society for the Promotion of Science (JSPS) (JP18H02370 and JP22K19300) and the Japan Agency for Medical Research and Development (AMED) (AMED PRIME, JP21gm6310017) to Y.M. Additional supports were provided by JSPS (20H03241 and 21H05720) and the Japan Science and Technology Agency (JST) FOREST (JPMJFR214L) to K.I, JST ACT X (JP1159335) to H.T, JSPS (JP21H02459) to Y.I., and MEXT Grants-in-Aid for Scientific Research (20H05926, 21K06053), and Grant from Institute for Fermentation, Osaka (G-2021-2-063) to S.C. N.U. was a recipient of the Support for Pioneering Research Initiated by the Next Generation (SPRING) program by JST (JPMJSP2157).

## Author contributions

N.U. and Y.M. designed the project. N.U. performed the experiments under the supervision by Y.M., K.I., H.T., and S.C. K.I. and Y.I. performed the mass spectrometry analysis. N.U. and Y.M. analyzed the data and drafted the manuscript. All the authors discussed the results and approved the manuscript for submission.

## Conflicts of interest

The authors declare no conflicts of interest associated with this manuscript.

## Notes

### Competing Interest Statement

The authors have declared no competing interest.

